# AggreBots: configuring CiliaBots through guided, modular tissue aggregation

**DOI:** 10.1101/2025.02.22.639695

**Authors:** D. Bhatttaram, K. Golestan, X. Zhang, S. Yang, Z. Gong, S. L. Brody, A. Horani, V. A. Webster-Wood, A. B. Farimani, X. Ren

**Affiliations:** Department of Biomedical Engineering, Carnegie Mellon University, Pittsburgh, USA; Department of Medicine, Washington University School of Medicine, St. Louis, USA; Department of Pediatrics, Washington University School of Medicine, St. Louis, USA; Department of Cell Biology and Physiology, Washington University School of Medicine, St. Louis, USA; Department of Mechanical Engineering, Carnegie Mellon University, Pittsburgh, USA

## Abstract

Ciliated biobots, or CiliaBots, are a class of engineered multicellular tissues that are capable of self-actuated motility propelled by the motile cilia located on their exterior surface. Correlations have been observed between CiliaBot motility patterns and their morphology and cilia distribution. However, precise control of these structural parameters to generate desired motility patterns predictably remains lacking. Here, we developed a novel Aggregated CiliaBot (AggreBot) platform capable of producing designer motility patterns through spatially controlled aggregation of epithelial spheroids made from human airway cells (referred to as CiliaBot Building Blocks or CBBs), yielding AggreBots with configurable geometry and distribution of active cilia. Guided multi-CBB aggregation led to the production of rod-, triangle-, and diamond-shaped AggreBots, which consistently effected greater motility than traditional single-spheroid CiliaBots. Furthermore, CBBs were found to maintain internal boundaries post-aggregation through the combined action of pathways controlling cellular fluidity and tissue polarity. This boundary fidelity, combined with the use of CBBs with immotile cilia due to mutations in the *CCDC39* gene, allowed for the generation of hybrid AggreBots with precision control over the coverage and distribution of active cilia, further empowering control of motility patterns. Our results demonstrate the potential of AggreBots as self-propelling biological tissues through the establishment of morphological “levers” by which alterations to tissue motility can be theoretically planned and experimentally verified.

## INTRODUCTION

Stem cells, with their inherent ability to replicate and differentiate in response to a wide variety of intrinsic and extrinsic cues, have long been seen as the quintessential building block for manufacturing *de novo* biological constructs. In particular, the ability of stem cells to self-assemble has been a key research area that has given rise to by-design functional outcomes (*1*). This controlled functionality has led to stem cells being of interest in the growing biohybrid and organic robotics field, where the ability of by-design function in living tissues is vital for controlling robot behavior (*2–7*). Biohybrid and organic robots, or biobots, are living constructs capable of self-actuation whereby their designs—both the composing cells and their organization—can drive motility, the ability to move independently powered by metabolic energy (*8–10*). The constituent cells within a biobot play a principal role in how the biobot actuates motility, with the most commonly investigated being muscle cells from cardiac or skeletal tissues, which produce forces through myofibril contraction (*4, 11–14*). However, myofibrils do not represent the only possibility for a force-generating building block when designing a biobot.

Beyond muscles, another possibility for generating macroscopic motility with living tissues lies in motile cilia, hair-like cellular appendages of approximately 200 nm in diameter ranging from 1-10 μm in length (*15, 16*). Each of these cilia are comprised of a carefully arranged collection of microtubules to which dynein motor proteins are bound that together produce a force-generating “effective stroke” followed by a “recovery stroke.” Cilia can be found on the apical surface of several epithelium-enclosed tissue compartments in the human body—such as the respiratory airways, brain ventricles, middle ears, and fallopian tubes (*17–24*). Coordinated cycles of effective and recovery strokes at a frequency of 10-40 Hz from a large number of cilia—up to trillions in the lungs—generate macro-scale motility in the form of metachronal waves that transport luminal substances, such as mucus-entrapped foreign particles from the lungs or eggs through the fallopian tubes to the uterus (*17, 18, 23, 24*). Looking back in evolution, motile cilia also represent an ancient mechanism for simple aquatic organisms to propel themselves through water environments, from unicellular eukaryotic ciliates like *Paramecium* that have relied on cilia-powered propulsion for hundreds of millions of years to the multicellular *Ctenophora* family of marine invertebrates that swim the world’s oceans via their cilia, being the largest animals to do so (up to 1.5 m in length) (*25, 26*).

Not until recently has the leap to human cell-derived ciliated biobots (CiliaBots) been made. These human CiliaBots are 3D organotypic mini-tissues (i.e., organoids) that are differentiated from stem cells of the human respiratory airways, thereby recapitulating the cellular composition of their tissue of origin, including the multiciliated cells (*1, 10, 27*). Prior work from us and others establishes that withdrawing extracellular matrix support from the culture environment is key to enabling the cilia-bearing apical epithelial surface to face the organoid’s outside. Such apical-out tissue polarity allows the exterior-facing cilia to drive macroscopic organoid motility, making them human CiliaBots. When embedded within a hydrogel substrate, these CiliaBots can generate stable motility in the form of 3D rotation, with the resulting activity being strongly linked to the beat frequency of constituent cilia (*27*). Intriguingly, comprehensive investigations of CiliaBot 2D locomotion when allowed to traverse a flat surface revealed a correlation between motility patterns and structural features such as tissue morphology and surface cilia coverage (*10*). But while these prior investigations provide valuable observations of spontaneously generated CiliaBot variations, a method for precisely *controlling* these structural parameters has yet to be established, presenting a challenge in developing designer CiliaBots with predefined structure and motility.

To address this need for manufacturing CiliaBots with predictable motility features by design, we present the concept and platform to engineer AggreBots. These higher-order CiliaBots leverage spatiotemporally controlled placement and aggregation of individual CiliaBot building blocks (CBBs). Each CBB is a cluster of a defined number of human airway stem cells, which on their own can differentiate into simple, spherical CiliaBots of reproducible size and high-density, homogeneous surface coverage of motile cilia. Aggregation of these reproducible CBBs in a spatially directed manner allows for control over resulting mature tissue geometry, which was utilized herein to generate several fundamental biobot designs, including rod-, triangle-, and diamond-shaped AggreBots. Additionally, we established a novel capability for control over the AggreBots’ active cilia coverage by introducing CBBs bearing *non-functional* cilia, leveraging airway stem cells with a genetic mutation responsible for primary ciliary dyskinesia (PCD) (*28*– *32*). The combination of controls over both tissue geometry and active cilia coverage, and by extension cilia distribution, allows for AggreBots to generate distinct and replicable patterns of 2D locomotion that have been herein quantified through their translational and rotational velocities. This analysis is further accompanied by a unique “path-and-extent” visualization method to transduce quantitative data into easily identifiable motility phenotypes (Figure 1). Taken together, the research herein solidifies the AggreBot as a next-generation CiliaBot platform with the potential to support applications in cell-based therapeutic delivery and ciliary theragnostics, due to the AggreBot platform’s cutting-edge design flexibility and reproducibility.

**Fig. 1.**
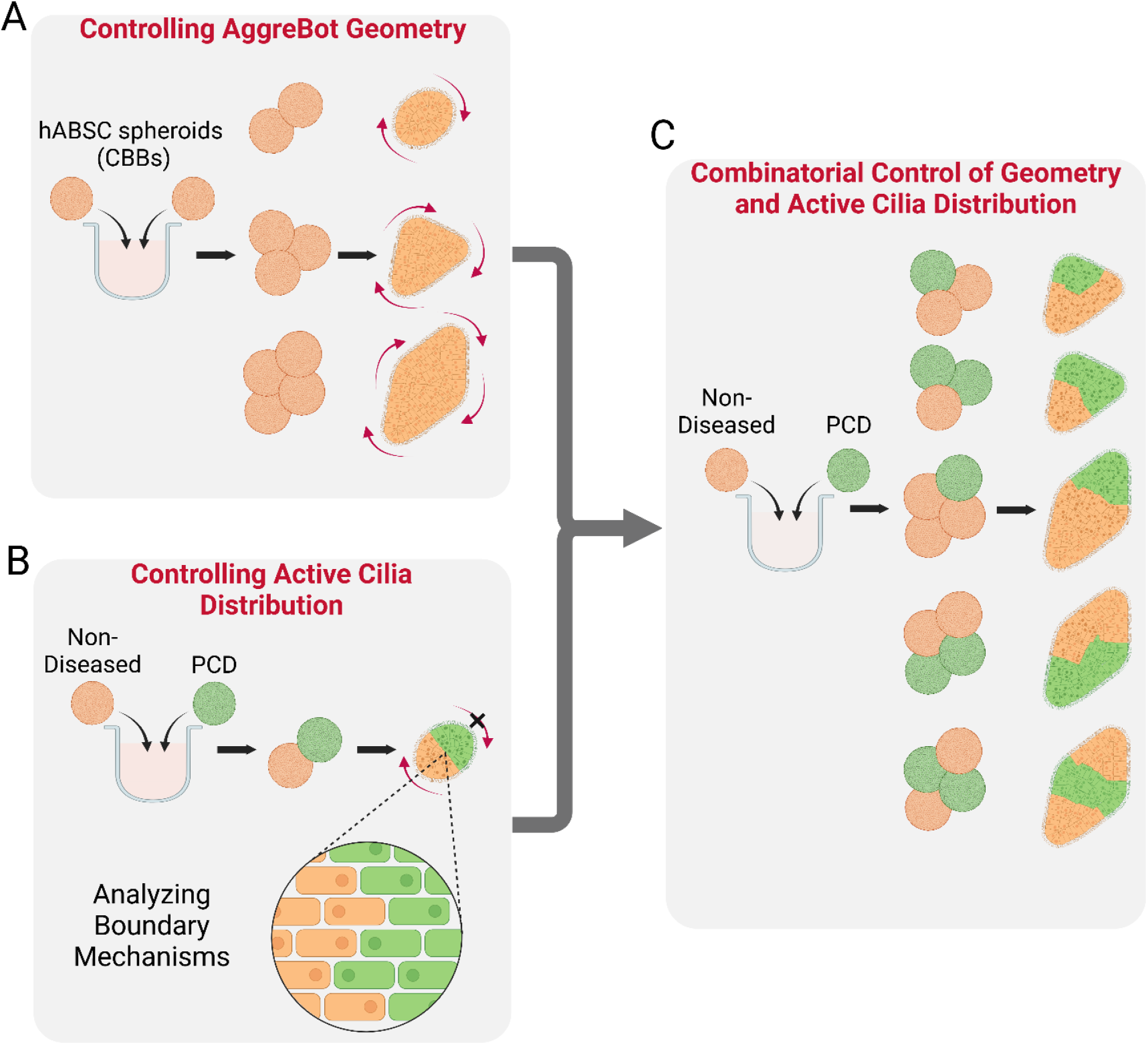
Schematic overview of AggreBot engineering. (**A**) Establishing geometric control of nascent AggreBots via spatiotemporally controlled aggregation of immature CBBs. (**B**) Establishing control of active cilia coverage and distribution in hybrid Aggrebots through the introduction of PCD-derived CBBs (to designate domains of cilia inactivity), along with investigating maintenance of the boundaries between constituent CBBs. (**C**) Utilizing combinatorial control of AggreBot geometry and active cilia distribution to generate ever-increasing permutations of AggreBots, effecting novel motility patterns. Full-color version available online.

## RESULTS

### Assembly of geometrically configurable CiliaBots through modular aggregation of CiliaBot Building Blocks (CBBs)

Our previously established CiliaBot protocol showcased the ability of human airway basal stem cells (hABSCs) isolated from non-diseased donors and seeded into a round-bottom, cell-repellent microplate to cluster into spherical tissues in the absence of ECM support. These tissues could then further differentiate into CiliaBots with an apical-out epithelial polarity where the exterior-facing motile cilia effectively propel tissue motility (*27*). However, this prior protocol did not allow for reproducible control of tissue geometry. Therefore, in this work, we first hypothesized that altering CiliaBot tissue geometry could be achieved through post-maturation aggregation— which could, by extension, have a controllable effect on CiliaBot motility. To explore this idea, we attempted to merge fully differentiated CiliaBots to generate CiliaBots with non-spherical geometries. However, this attempt failed due to interference from the constant cilia beating on the CiliaBots’ exterior surfaces that repelled one CiliaBot from the other, preventing stable contact (Figure S1, Movie S1).

Based on these initial findings, we subsequently assessed the effect of aggregation of epithelial spheroids prior to tissue maturation and cilia emergence. Seeding of a set number (500) of non-diseased hABSCs derived from normal tissue in a cell-repellent, U-bottom microwell surface, formed spheroids with homogeneous size distribution. These 1-day-old spheroids were then paired and reseeded into the same well, where pairs were brought into contact by U-bottom well curvature. Within as soon as 2 hours following initial pairing, adhesion and aggregation of the spheroid pairs was readily observed, compacting in length and expanding in width as culture continued (Figure 2A).

**Fig. 2.**
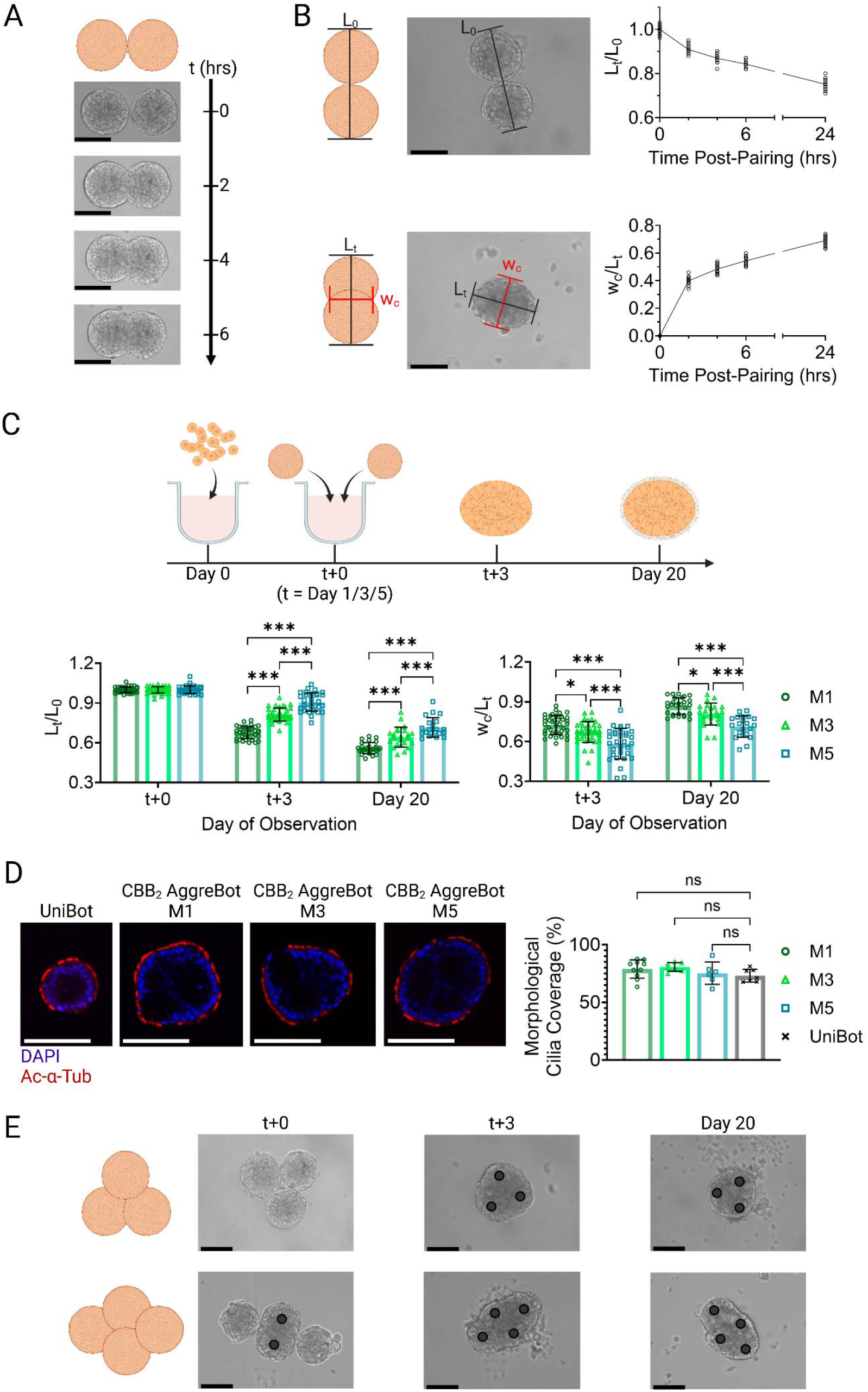
Characterization of CBB aggregation and resulting AggreBot morphology. Time series images of aggregation behavior of 2 CBBs between 0 and 6 hours after pairing. (**B**) Depiction and quantification of normalized length (L_t_/L_0_) and width-length ratio (w_c_/L_t_) over the course of CBB_2_ aggregation of 1-day-old CBBs between 0 and 24 hours after pairing, with accompanying schematic. (**C**) Quantification of normalized length (L_t_/L_0_) and width-length ratio (w_c_/L_t_) of 2-CBB aggregation of CBBs aged 1, 3, and 5 days. Quantification was done immediately following CBB pairing, 3 days following pairing, and as the AggreBot approached tissue maturation at Day 20, with accompanying schematic. (**D**) Example confocal slices of UniBots and CBB_2_ AggreBots with immunostaining of Ac-α-Tub and DAPI, along with quantification of morphological cilia coverage. (**E**) Time series images of CBB_3_ triangle-shaped and CBB_4_ diamond-shaped AggreBot formation with images shown immediately following pairing with last CBBs introduced, 3 days following pairing, and immediately prior to tissue maturation at Day 20. Black circle markers overlayed on images to aid in identification of constituent CBBs. All data represent means ± SD. *: *p* < 0.05, **: *p* < 0.01, and ***: *p* < 0.001. Two-way ANOVA with Tukey’s multiple comparisons test in panel (**C**) and one-way ANOVA with Tukey’s multiple comparisons test in panel (**D**). Scale bar in all panels, 125 μm. Full-color version available online.

To quantify the extent of spheroid aggregation observed in the paired culture process, two morphological metrics were developed: (1) normalized length, defined as the length of the aggregate’s major axis (L_t_) as a ratio of the average initial pre-aggregation end-to-end lengths (L_0_) of the spheroid pairs; and (2) width-length ratio, defined as the ratio between the aggregate’s central width (w_c_, corresponding to the minor axis) and major axis length. With these metrics, we observed highly regular lengthwise compaction of two spheroids (with normalized length reaching 0.75±0.02 (mean ± std at 24 hours post-pairing) and central width expansion (width-length ratio of 0.69±0.03 (mean ± std at 24 hours post-pairing) in aggregated spheroids (Figure 2B). These findings indicate that immature hABSC spheroids, which do not yet present functional cilia on their exterior surface, could be used as CiliaBot building blocks (CBBs) capable of aggregating in controlled geometries. Based on these findings, we theorized that these aggregates can then be matured into CiliaBots of any desired geometry (AggreBots), in contrast with the spherical CiliaBots derived from one CBB (UniBots).

To generate more longitudinal observations of AggreBot morphology, we maintained rod-shaped AggreBots derived from 2 CBBs (denoted CBB_2_) up until tissue maturation (at minimum 21 days post seeding), imaging the aggregate at initial pairing (t+0), 3 days after initial pairing (t+3), and at Day 20 post *seeding* as the AggreBots approached maturity. Furthermore, to evaluate for any alterations to the aggregation behavior resulting from the age of the CBB (number of days following hABSC seeding for CBB formation) at the time of initial pairing, we included in this test CBBs that were 1, 3, and 5 days old prior to being paired (the abbreviations M1, M3, and M5 are used to denote CBB age at the timepoint of first pairing and subsequent <u>m</u>erging). We observed the same tendency towards lengthwise compaction and central width expansion with these experimental conditions as seen in the initial paired seeding experiments (Figure 2C). However, the bulk of tissue reorganization occurs in the early days after pairing (i.e., normalized length decreasing from 1.00±0.02 to 0.68±0.04 (mean ± std) after the first 3 days of aggregation but decreasing further to only 0.56±0.04 (mean ± std) in the 16 days thereafter). Additionally, significant differences in both the normalized lengths (*p* < 0.001) and width-length ratios (*p* < 0.05) of the AggreBots stemming from CBB pairing at differing times were observed, with older CBB pairings tending to aggregate less in both length and width (Figure 2C). Consistent with this, we observed a decrease in aggregation success rate corresponding to increased age of CBB (Figure S2), indicating that pairing of CBBs at early timepoints is critical to the successful formation of AggreBots.

To assess any effects of aggregation on ciliogenesis and cilia coverage on the exterior tissue surface, mature AggreBots were evaluated with cilia marker Acetylated-α-Tubulin (Ac-α-Tub) (Figure 2D). Ciliation of the AggreBots’ outer surface was then calculated from the middle slices of the AggreBots’ confocal z-stacks, measured as a percentage of the AggreBot surface covered in Ac-α-Tub signal (*33*). Staining of un-aggregated CBBs that subsequently differentiated into UniBots was conducted as a control comparison. We found no significant difference in surface cilia coverage between the AggreBots of any pre-pairing CBB age when compared to UniBots (*p* > 0.05), indicating the CBB aggregation process and increased CiliaBot dimension from UniBot to AggreBot does not interfere with multiciliated cell differentiation and apical cilia presentation (Figure 2D).

After establishing the foundation of modular CBB aggregation with pairs delivering rod-shaped AggreBots at the time of tissue maturation, we pursued geometric configurations of increased complexity with CBB_3_ triangle-shaped and CBB_4_ diamond-shaped protocols, relying on the same well curvature as in the rod-shaped CBB_2_ studies to bring the CBBs into contact (Figure 2E). Following the maturation period, triangle- and diamond-shaped AggreBots were achieved, largely retaining the shape of the initial geometric configuration.

### Regulation of AggreBot motility by tissue geometric configuration

Following from our findings that CBB aggregation did not significantly impact surface cilia coverage, we theorized that an AggreBot would exhibit increased motility compared to a UniBot of equal cell count. The AggreBots, we reasoned, would have a larger surface area and thus an increased number of exterior-facing cilia for tissue propulsion while possessing near-equal volume to the UniBot. We further hypothesized, circling back to our overarching aim, that the control over CiliaBot tissue geometry aggregation provides would entail control over resulting motility. To test for these hypotheses, we introduced the biomechanical assay of a 2D locomotion test, where mature CiliaBots would be allowed to move free on a flat-bottom microplate (Figure S3) (*10*).

Prior to addressing these hypotheses, we sought to develop metrics for measuring CiliaBot motility. The majority of CiliaBots observed displayed a progressing spiral motion akin to a loop-de-loop (Movies S2-4). These observations motivated the extraction of two primary motility metrics for CiliaBots: (1) the average translational speed, denoted v, representing the change in XY-position of the CiliaBot centroid over the video recording time; and (2) the average rotational speed, denoted ω, representing the change in the orientation of the CiliaBot’s major axis over the same period (Figure 3A). From these, the metric of average path curvature, denoted κ, was calculated, derived from the ratio of the two previous metrics (ω/v), representing the tightness of the spiral motion the CiliaBot generated and, by extension, a quantification of its motility pattern. In addition to the development of these quantitative metrics, a visualization method of CiliaBot motility patterns was designed known as the “path-and-extent” model, where the curve of the CiliaBot centroid is traced, denoting the path of locomotion, and the CiliaBot’s vertices, taken as a representation of the full space the CiliaBot traverses, represent the extent (Figure 3B). The result is a visualization method applicable to both AggreBots and UniBots, delivering a distillation of any given motility pattern (Figure 3C).

**Fig. 3.**
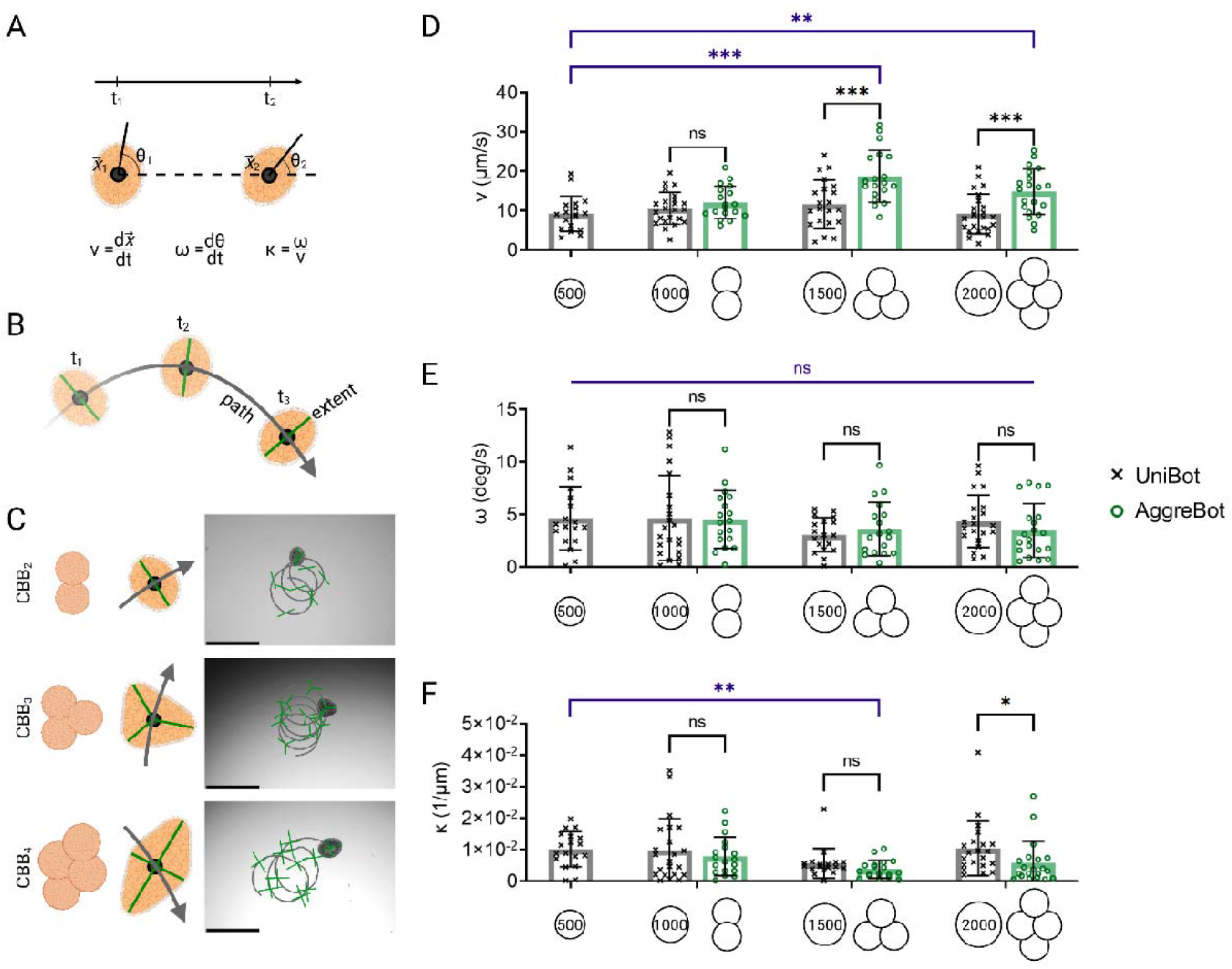
Characterization of AggreBot motility in comparison to UniBot motility. (**A**) Schematic depiction of quantification of translation velocity (v), rotational velocity (ω), and path curvature (κ) through analysis of CiliaBot position (x) and angle of orientation (θ). (**B**) Schematic depiction of “path-and-extent” visualization method developed for displaying both CiliaBot position and angle of orientation over time. (**C**) Example “path-and-extent” depictions of CBB_2_, CBB_3_, and CBB_4_ AggreBot motility, with associated schematics indicating the “vertebral” positioning for each AggreBot type. Scale bar, 500 μm. (**D**-**F**) Quantifications of CiliaBot translational velocity (**D**), rotational velocity (**E**), and path curvature (**F**) for CBB_2_, CBB_3_, and CBB_4_ AggreBots in comparison with UniBots of corresponding cell count. All data represent means ± SD. *: *p* < 0.05, **: *p* < 0.01, and ***: *p* < 0.001. Two-way ANOVA with Tukey’s multiple comparisons test applied to the 1000-, 1500-, and 2000-cell CiliaBot groups (black) and one-way ANOVA with Tukey’s multiple comparisons test applied to the 500-cell UniBot group and all AggreBot groups (blue) in panels (**D**-**F**). Full-color version available online.

To test the hypothesis of increased surface area having a positive effect on motility, we stratified the overarching classifications of AggreBot and UniBot based on their hABSC counts at formation, ranging from 1000 to 2000 cells, from which any difference in motility between an AggreBot and a UniBot in a given cell “weight class” could only be reasonably attributed to tissue configuration and the resulting surface area and cilia quantity. An analysis of the motility of the 500-cell UniBot was further included in this setup to test the hypothesis of tissue geometry impacting the patterns of CiliaBot motility, with a separate statistical test running in parallel comparing the 500-cell UniBot to the 3 tested AggreBot forms. A significant increase in average translational speed in 1500-cell CBB_3_ and 2000-cell CBB_4_ AggreBots compared to their 1500- and 2000-cell UniBot counterparts was observed (*p* < 0.001) (Figure 3D); in contrast, the speeds of 1000-cell CBB_2_ AggreBots were not significantly different than their 1000-cell UniBot counterparts (*p* = 0.37). Across all tested cell counts, no significant differences were observed in average rotational speed (*p* > 0.05), an observation that extended to average path curvature, in turn, with the exception of the 2000-cell class (*p* = 0.04) (Figure 3E, 3F). When comparing the motility of the various AggreBot classes to the 500-Cell UniBot, the translational speed of the 1500-cell CBB_3_ (*p* < 0.001) and 2000-cell CBB_4_ (*p =* 0.009) AggreBots were found to be significantly elevated (Figure 3D). No significant differences were observed in rotational speed (*p* > 0.05), while the path curvature of the 1500-cell CBB_3_ AggreBot was found to be significantly increased (*p =* 0.006) (Figure 3E, 3F). This two-part experiment established that aggregation of CBBs into AggreBots can produce additional motility in excess of non-aggregated CiliaBots of the same size, while also establishing the manipulation of CiliaBot geometry as a pathway towards control over motility patterns, though tissue configuration represents only one lever of control.

### Controlling active cilia coverage leveraging immotile-cilia CBBs

With control over tissue geometry established, we determined that control over the surface coverage of active cilia would provide us with an additional “lever” with which to direct CiliaBot motility. In particular, we theorized that the ability to generate discrete “active” and “inactive” zones on the tissue surface would further improve motility control. To this end, we acquired hABSCs bearing genetic mutations in the *CCDC39* gene that cause primary ciliary dyskinesia (PCD), a disorder that can affect both the structure and function of cilia—in the case of the *CCDC39* mutation, immotile cilia are produced (*28–32, 34, 35*). We observed that solitary CBBs formed from PCD hABSCs (denoted as ^PCD^CBBs), underwent the same tissue reorganization and maturation seen in non-diseased hABSC-derived CBBs (denoted as ^ND^CBBs), ultimately producing “cilia-dead” UniBots that displayed limited motility when compared to “cilia-active” UniBots from ^ND^CBBs (Figure 4A, Movie S5,6), findings corroborated by our previous work (*27*).

**Fig. 4.**
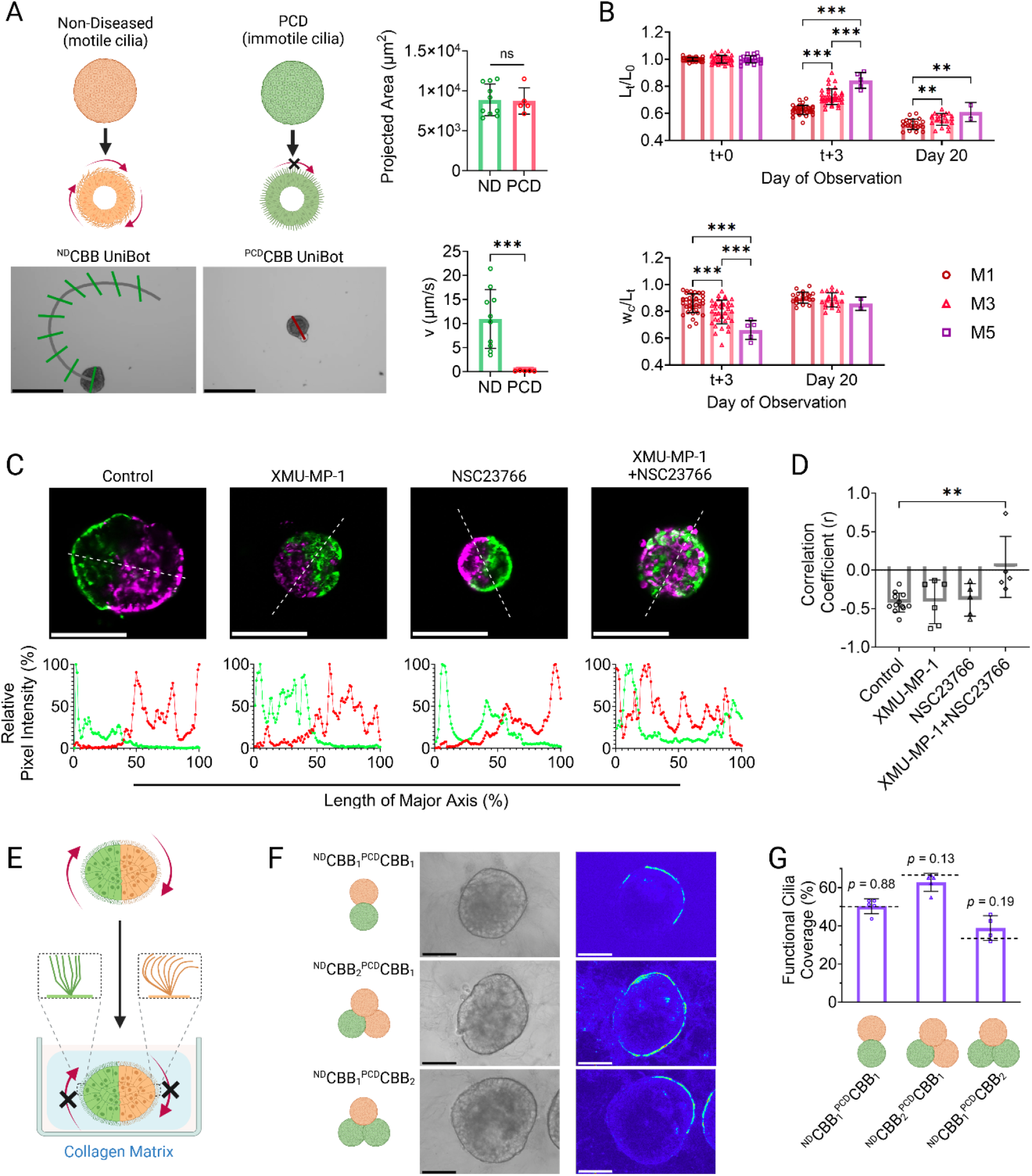
Incorporation of ^PCD^CBBs to generate hybrid AggreBots to control active cilia coverage. (**A**) Example “path-and-extent” depictions of ^ND^CBB UniBot and ^PCD^CBB UniBot motility, along with quantitative comparisons of morphology and motility between the UniBot classes. Scale bar, 250 μm. (**B**) Quantification of normalized length (L_t_/L_0_) and width-length ratio (w_c_/L_t_) of hybrid ^ND^CBB_1_^PCD^CBB_1_ aggregation at CBB ages of 1, 3, and 5 days. Quantification was done immediately following CBB pairing, 3 days following pairing, and as the AggreBot approached tissue maturation at Day 20. (**C**) Example confocal slices of fluorescently dyed ^ND^CBB_2_ AggreBot treated with 10 μM YAP activator XMU-MP-1, 50 μM Rac1i inhibitor NSC23766, both in combination, or neither, along with example quantifications of relative pixel intensity for DiO (green) and DiD (red) signal across the length of the AggreBot’s major axis as determined by an image profile. Scale bar, 125 μm. (**D**) Quantification of correlation coefficients between paired green and red relative intensities across AggreBot image profile when treated with XMU-MP-1, NSC23766, both chemicals, or neither. (**E**) Schematic depiction of collagen embedding of AggreBots to arrest tissue-level motility and allow for analysis of functional cilia coverage in hybrid AggreBots. (**F**) Example depictions of collagen-embedded hybrid AggreBots, along with associated heatmaps of pixel intensity variation over the course of videotaping. (**G**) Quantification of functional cilia coverage in hybrid AggreBots, with associated *p-* values for one-sample t-tests comparing observed coverage to expected values (black dashed lines). Scale bar, 75 μm. All data represent means ± SD. *: *p* < 0.05, **: *p* < 0.01, and ***: *p* < 0.001. Unpaired t-test in panel (**A**), two-way ANOVA with Tukey’s multiple comparisons test in panel (**B**), one-way ANOVA with Tukey’s multiple comparisons test in panel (**D)**, and one-sample t tests in panel (**G**). Full-color version available online.

To determine whether ^PCD^CBBs could be combined with ^ND^CBBs for AggreBot generation, we applied the same aggregation protocol previously utilized for ^ND^CBB_2_ to the new hybrid (denoted ^ND^CBB_1_ ^PCD^CBB_1_). We observed similar aggregation dynamics in the hybrid to ^ND^CBB_2_ counterparts, including the bulk of lengthwise compaction and central width expansion taking place in the early days of aggregation. Later aggregating CBBs also produced more elongated and thinner AggreBots like in the ^ND^CBB_2_ experiment, though by Day 20, the differences in width-length ratio were not found to be statistically significant (*p* > 0.05) (Figure 4B).

Upon demonstrating effective ^PCD^CBB-to-^ND^CBB aggregation, we assessed whether CBBs merged within an AggreBot maintain distinct domains within the larger structure, which, if true, would be the key to achieving spatially controlled functional cilia coverage via guided placement of ^PCD^CBBs and ^ND^CBBs. To this end, we undertook investigations to assess both the structural and functional distinctiveness of CBB domains within AggreBots. Rod-shaped ^ND^CBB_2_ AggreBots were generated, one labeled with DiO fluorescent dye (green) and one labeled with DiD (red). Intriguingly, despite being derived from the same hABSC source and cultured under the same conditions, the labeled CBBs, while fusing into an AggreBot, did not intermix their constituent cells, instead maintaining a clear boundary between themselves (Figure 4C).

Regulators of both cellular fluidity and tissue polarity are known to regulate cell migration in epithelial tissue development (*36–39*). To mechanistically investigate their involvement in boundary establishment between CBB domains within AggreBots, we treated these dual-color CBB aggregates with XMU-MP-1, an activator of YES-associated protein (YAP) known to increase the fluidity of lung epithelial cells through the inhibition of a process known as “jamming”, and with NSC23766, an inhibitor of Rac1 GTPase, a protein responsible for the establishment of apical polarity in epithelial tissues. While neither small molecule alone disrupted the inter-CBB boundary, their combination led to the intermixing of cells with distinct fluorescence labeling, as evidenced by the loss of negative correlation between regions of high green signal and high red signal in the combination group, suggesting boundary disruption (Figure 4D).

With the structural distinctiveness of post-aggregation CBBs established, we carried hybrid ^ND^CBB-^PCD^CBB AggreBots to maturity to assess the functional distinctiveness of cilia activity. To this end, rod- and triangle-shaped hybrid AggreBots were generated and immobilized in collagen matrix to limit their 2D locomotion while allowing for close observation of surface cilia beating (Figure 4E). In this environment, we observed distinct domains of cilia activity and inactivity, matching the configuration of ^ND^CBB and ^PCD^CBB as initially aggregated (Figure 4F, Movie S7-9). Quantitative analysis of the embedded AggreBots further corroborated these findings (Figure 4G), as the calculated functional cilia coverage data for the ^ND^CBB_1_ ^PCD^CBB_1_, ^ND^CBB_2_ ^PCD^CBB_1_, and ^ND^CBB_1_ ^PCD^CBB_2_ groups were not found to be significantly different from the theoretical coverage based on the AggreBots’ compositions (50%, 66.7%, and 33.3%) (*p* > 0.05). This, together with the results of fluorescent CBB domain tracking (Figure 4D), established the feasibility of using ^PCD^CBBs for fine control of active cilia coverage on the AggreBot surface.

### Combinatorial control of geometry and active cilia coverage to effect novel motility patterns

With control over both tissue geometry and active cilia coverage established, we sought to quantify the effects of combinatorial manipulation of these independent variables and the effects this might have on AggreBot motility. To this end, we constructed rod-, triangle-, and diamond-shape AggreBots with either all ^ND^CBBs or a combination of ^ND^CBBs and ^PCD^CBBs and compared their 2D locomotion. To assess and visualize motility, the “path-and-extent” visualization method was modified to allow clear distinction of AggreBot types via green vs. red centroid-vertex segments—corresponding to researcher input—indicating ^ND^CBB vs. ^PCD^CBB domains, respectively (Figure 5A i-vi, Movie S10-15).

**Fig. 5.**
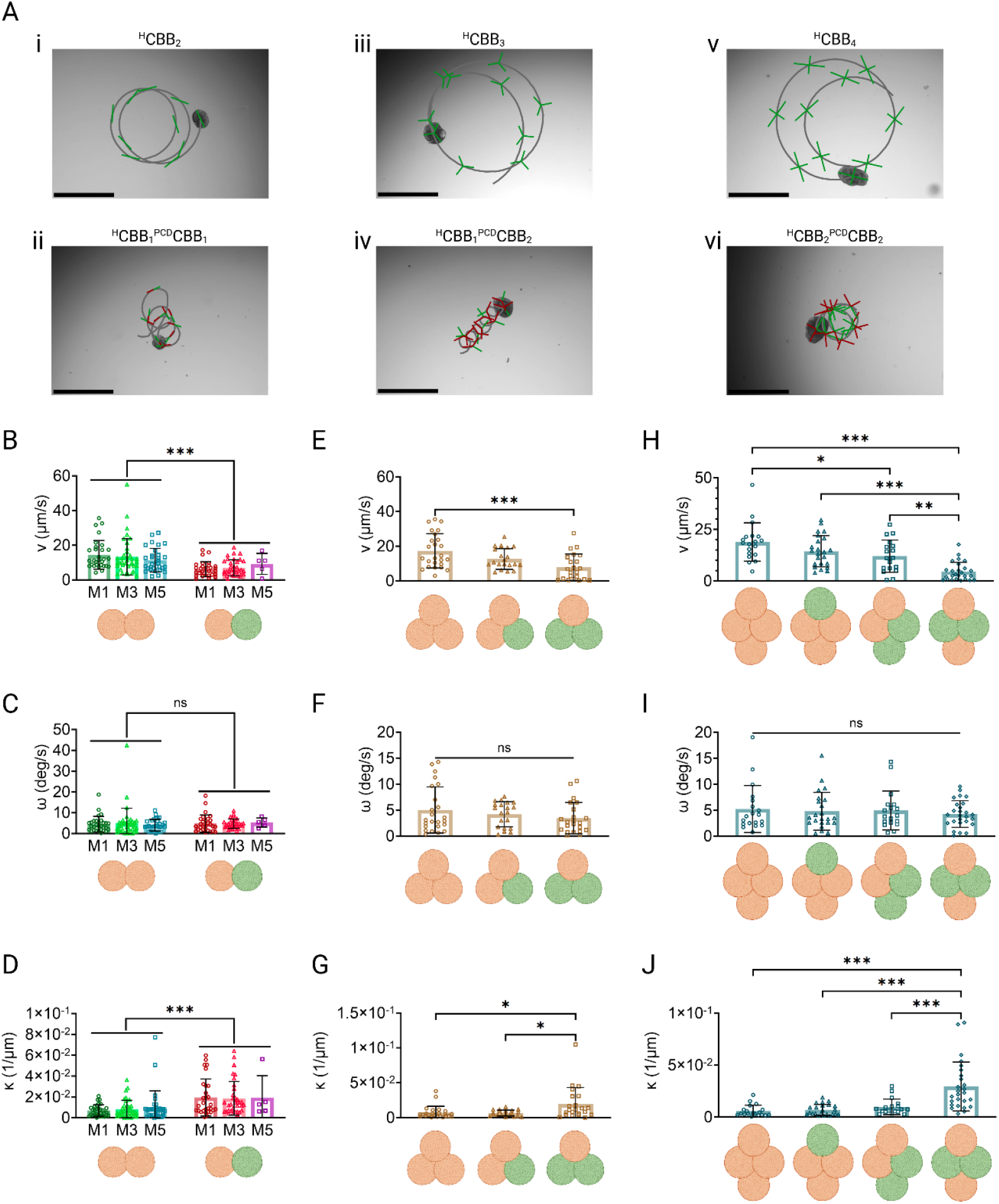
Characterization of hybrid AggreBot motility in comparison with all-^ND^CBB AggreBots. (**A**) Example “path-and-extent” depictions of all-^ND^CBB and hybrid CBB_2_, CBB_3_, and CBB_4_ AggreBots. Scale bar, 500 μm. (**B**-**D**) Quantifications of CBB_2_ AggreBot translational velocity (**B**), rotational velocity (**C**), and path curvature (**D**), comparing all-^ND^CBB and hybrid AggreBots aggregated with 1-, 3-, and 5-day-old CBBs. Orange spheroids represent ^ND^CBBs while green spheroids represent ^PCD^CBBs. (**E**-**G**) Quantifications of CBB_3_ AggreBot translational velocity (**E**), rotational velocity (**F**), and path curvature (**G**), comparing all-^ND^CBB and hybrid (^ND^CBB_2_^PCD^CBB_1_ and ^ND^CBB_1_^PCD^CBB_2_) AggreBots. Orange spheroids represent ^ND^CBBs while green spheroids represent ^PCD^CBBs. (**H**-**J**) Quantifications of CBB_4_ AggreBot translational velocity (**H**), rotational velocity (**I**), and path curvature (**J**), comparing all-^ND^CBB and hybrid (^ND^CBB_3_ ^PCD^CBB_1_ chiral ^ND^CBB_2_ ^PCD^CBB_2_, and symmetric ^ND^CBB_2_ ^PCD^CBB) AggreBots. Orange spheroids represent ^ND^CBBs while green spheroids represent ^PCD^CBBs. All data represent means ± SD. *: *p* < 0.05, **: *p* < 0.01, and ***: *p* < 0.001. Two-way ANOVA with Tukey’s multiple comparisons test in panels (**B**-**D**) and one-way ANOVA with Tukey’s multiple comparisons test in panels (**E**-**J**). Full-color version available online.

Starting with the rod-shaped Aggrebots, we observed a significant decrease in average translational speed in the ^ND^CBB_1_ ^PCD^CBB_1_ configuration compared to the ^ND^CBB_2_ (*p* < 0.001), which proved indicative of a trend of ^PCD^CBB introduction leading to a decrease in translational speed (and an accompanying increase in average path curvature). This trend is also seen in triangle-shaped AggreBots, where ^ND^CBB ^PCD^CBB_2_ showcase significantly reduced translational speed (*p* < 0.001) and increased path curvature (*p =* 0.02) when compared to ^ND^CBB_3_ AggreBots, with ^ND^CBB_2_ ^PCD^CBB_1_ AggreBots exhibiting motility patterns between these extremes. Of particular note are the differing motility results within the tested diamond-shaped AggreBots. While the same trend seen in rod- and triangle-shaped AggreBots was present, the increased number of constituent CBBs per diamond-shaped AggreBot allows for a higher number of permutations in which the ^ND^CBB_2_ and ^PCD^CBB_2_ could be arranged, leading to two separate tested diamond AggreBot configurations with an overall composition of ^ND^CBB ^PCD^CBB (Figure 5H-J). Here, we found that the different permutations (in particular, the locations of the ^ND^CBB within the larger structure) resulted in significantly altered translational speed (*p =* 0.007) and path curvature (*p* < 0.001). The chiral ^ND^CBB_2_ ^PCD^CBB_2_ variant was more likely to adopt a loop-de-loop locomotion pattern. In contrast, the symmetric variant instead displayed minimal translational speeds but substantial rotational speeds, indicating rotation in place (Figure 5A, 5H-J).

## DISCUSSION

In this work, we established the AggreBot as a cutting-edge platform for generating designer CiliaBots with defined morphology and active cilia coverage. To accomplish this, we established a modular approach to spatially control the aggregation of CBBs to serve as building blocks for higher-order CiliaBots; this was accomplished while maintaining homogeneous coverage of motile cilia on the CiliaBot’s exterior surface upon tissue maturation. We further established control over the extent of surface active cilia coverage, incorporating PCD-derived, “cilia-dead” CBBs into the aggregation protocol to deliver designer cilia distribution and novel motility patterns.

The research herein provides a comprehensive understanding of the aggregation behavior of epithelial spheroids, specifically multicellular clusters of epithelial stem cells derived from the human airways. Epithelial spheroid aggregation is a phenomenon that was reported on previously in CiliaBot research (*10*). In making this process the focal point of our work, we have found spheroid aggregation to be highly reproducible and characterizable, allowing us to additionally tease out the time-dependent nature of this aggregation process. The discovery that both the success rate and the extent of aggregation can be influenced by the age of CBBs prior to aggregation indicates to us the possibility that the cellular composition of CBBs drives epithelial aggregation behavior as they progressively differentiate from a stem cell state toward their multiciliated cell fate. The CBB aggregation platform and our associated morphological characterization pipeline and preliminary findings could serve as the basis for future research in this discipline, which we anticipate may further expand our abilities to control the manner and extent of CBB aggregation and, ultimately AggreBot geometry.

The long-term maintenance of boundaries between constituent CBB domains post-aggregation is another fascinating advancement in our understanding of epithelial merging, in addition to the nature of the studied chemical effectors that then disrupted this boundary maintenance. Activation of the YAP-Taz signaling pathway has been shown to encourage epithelial cell fluidity, while inhibition of Rac1 hampers the establishment of apical-basolateral polarity (*36–39*). The finding that the combination of these effectors ultimately disrupted the inter-CBB boundary (with either chemical alone proving insufficient) suggests multiple inherent cellular mechanisms function in a redundant manner to prevent epithelial migration across predefined tissue domains, which further speaks to the robust nature of our AggreBots maintaining distinct domains throughout maturation.

Furthermore, this work pioneered controlling CiliaBot geometry—and, by extension, motility— through the modular assembly of uniform CBBs. The regularity of the underlying CBBs, in terms of cellular composition, tissue dimension, differentiation potential, and aggregation behavior, is crucial to the AggreBot platform introduced in this research, without which shape planning and resulting aggregation protocols would not have been possible. Both the current and prior investigations have demonstrated that overnight culture of hABSCs under cell-repellent conditions in the absence of ECM support is a simple yet effective way to generate CBBs with these desired features (*1, 10, 27*). Additionally, further mimicking the homogeneity previously reported in mature UniBots, we demonstrate aggregation behavior without sacrificing uniform cilia coverage on the tissue’s outer surface (*27*). Together, these pillars allow for the delivery of high-level shape control in mature AggreBots, resulting in enhanced translational motility over UniBots with the same cell count. This increased power-to-weight ratio is especially promising when considering the scalability of CiliaBot motility; in designing CiliaBots of greater size in the future, aggregation of a higher number of CBBs may prove crucial, though additional external guides to achieve increased complexity, such as guiding molds or bioprinting, may be required (*40–43*). A further consideration is that as AggreBot size and complexity increase with the introduction of supplementing technologies, future research in this discipline may be confronted with ethical concerns, potentially regarding whether such constructs constitute living entities. This question could be further compounded due to the use of human cell lines in the AggreBot’s construction. Going forward, such challenges will need to be addressed in order to ensure ethical compliance (*44*).

Prior reports showed that CiliaBots could be generated with spontaneous variations in morphology (i.e., spherical vs. elongated) and surface cilia (i.e., fully covered vs. polarized), which could, in turn, be correlated with specific motility patterns (*10*). These findings propelled us to not only observe active cilia coverage, but also to directly control it through the novel use of ^PCD^CBBs made from PCD cells bearing genetic mutations resulting in non-functional cilia to designate “cilia-dead” domains. Through the geometrically controlled aggregation of ^PCD^CBBs with the “cilia-active” ^ND^CBBs, we demonstrated hybrid AggreBots with robust control over-active cilia coverage. This novel type of CiliaBot was made possible through the combination of the following foundational observations. First, CBBs made from non-diseased and PCD cells behaved similarly in terms of CBB formation and subsequent differentiation towards multiciliated cells, resulting in tissue domains that are structurally near-indistinguishable but deliver significantly different functional outputs, a finding corroborated by previous research (*27*). Second, CBBs with distinct genetic backgrounds merged effectively, displaying similar aggregation behavior to CBBs with the same genotype. Lastly, CBBs showcased an inherent ability to maintain an internal boundary post-aggregation. These three discoveries allow mature AggreBots of both homogeneous and hybrid compositions to faithfully represent their initial designs and predictably diversify CiliaBot motility patterns. Our efforts in AggreBot development were further supported by our newly established analytical pipeline to capture CiliaBot motility as a combination of translation and rotation. We anticipate that this visual distillation of AggreBot behavior will allow for the rapid identification of desirable CBB compositional groupings in the future.

Finally, the key finding that CiliaBot motility can be varied not only through the underlying CBB composition of the AggreBot but also through the permutation of CBB positions within the AggreBot, as evidenced by the differences in the motilities of symmetric and chiral ^ND^CBB_2_ ^PCD^CBB_2_ AggreBots, is perhaps the most captivating of our results. This finding further underscores the controllability of the AggreBot platform when designing higher-order CiliaBots. Simply by varying the order in which the CBBs were aggregated, new patterns of locomotion were identified and replicated. This finding highlights a third lever of structural control in CiliaBot generation that directly emerges out of the control over geometry and cellular composition. With this modulation of CBB permutation, we achieve full control of active cilia distribution along with overall coverage. Through these levers provided by the AggreBot platform, we open the door to constructing living propeller systems with pre-planned, patterned ciliation, rather than the ubiquitous ciliation of UniBots, serving as a key step forward in controlling CiliaBot motility. Furthermore, this research has established the foundation for the modular design and assembly of scalable CiliaBots, having progressed from an axial length of 100 μm to 500 μm over the span of this research alone. The tools, techniques, and findings of this work, therefore, substantially advance the capabilities and reproducibility of ciliated biobots, enabling future studies to move beyond the bespoke design of biobots for individualized purposes toward true engineered design of robust, motile CiliaBots for specific applications in biomedical research, treatment, and theragnostics.

## MATERIALS AND METHODS

### Culture of human airway basal stem cells (hABSCs)

hABSCs from non-diseased donors were obtained from Lonza (Cat# CC-2541). hABSCs carrying primary ciliary dyskinesia (PCD)-associated genetic mutations in the CCDC39 gene (c.830_831delCA (p.Thr277Argfs*3) and c.1871_1872delTA (p.Ile624Lysfs*3)), leading to immotile cilia, were isolated from de-identified tissues with permission from the institutional review board at Washington University in Saint Louis (IRB# 201705095). Both non-diseased- and PCD-hABSCs (P3-5) were maintained at 37°C with 5% CO_2_ in Bronchial Epithelial Cell Growth Medium (Lonza, CC-3171) supplemented with 1 μM A83-01 (Sigma-Aldrich, SML0788), 5 μM Y-27632 (Cayman Chemical, 129830-38-2), 0.2 μM of DMH-1 (Tocris, 4126), and 0.5 μM of CHIR99021 (REPROCELL, 04000402) on T-25 flasks (Greiner Bio-One, 690175) pre-coated with conditioned medium from 804G cells, with initial plating density of 3500 cells/cm^2^ (*27, 45*). A full media change was conducted every 2 days after plating, with daily observations of cell confluency within the flask.

### Generation of CiliaBot Building Block (CBB) and differentiation into UniBot

hABSCs (P3-5) between 50 and 60% conflucency were lifted from T-25 flasks with TrypLE (ThermoFisher Scientific, 12563011) and resuspended in differentiation medium (PneumaCult-ALI Medium (STEMCELL Technologies, 07925) supplemented with 1 μM A83-01 and 5 μM Y-27632). Single-cell suspension of hABSCs were then seeded into U-bottom 96-well cell-repellent microplates (Greiner Bio-One, 655970) at the desired number of cells (500, 1000, 1500, 2000, or 3000) per well, and allowed to cluster overnight. The resulting hABSC cell spheroid was referred to as a CBB. CBBs derived from non-diseased- and PCD-hABSCs were referred to as ^ND^CBB and ^PCD^CBB, respectively. To produce a spherical CiliaBot consisting of a single CBB (referred to as UniBot), each solitary CBB was further differentiated for 21-28 days with half media changes every other day and incubated at 37°C with 5% CO_2_ to reach full maturity (*27*).

### Aggregation of CBBs and differentiation into AggreBot

To engineer Aggrebots, defined numbers of 1-day-old CBBs were placed in the same well within a U-bottom 96-well cell-repellent microplate in a geometrically controlled manner, and allowed to aggregate and further differentiate for 21-28 days to reach full maturity under the conditions described above for CBBs. Rod- and triangle-shaped AggreBots were generated through controlled aggregation of two and three CBBs, respectively. For diamond-shaped AggreBots, to enable consistent placement of CBBs during aggregation, two 1-day-old CBBs were allowed to aggregate first overnight, followed by the introduction of two additional, 2-day-old CBBs. Aggrebots were produced either all from ^ND^CBBs, or from a combination of ^ND^CBB(s) and ^PCD^CBB(s) as specified in the Results section. To investigate how CBB age influenced aggregation behavior, rod-shaped AggreBots were also aggregated from two CBBs on day-3 and - 5 of differentiation.

### Morphological analysis of rod-shaped AggreBots

Rod-shaped AggreBots derived from two CBBs were imaged under brightfield utilizing an EVOS M7000 microscope (ThermoFisher Scientific) immediately after CBB pairing, 3 days after aggregation, and as the AggreBot approached maturity (20 days after initial CBB formation). The 2D projections of the AggreBots from these images were manually traced and analyzed using custom MATLAB programs that extracted the major axis of the projection (representing the AggreBot length) and the bisecting line of the projection’s area running perpendicular to the major axis (representing the AggreBot width) at each of the timepoints of imaging, which were further used to calculate the normalized length and width-length ratio.

### CiliaBot motility analysis and visualization

For analysis and visualization, mature AggreBots and UniBots (collectively CiliaBots) aged 21-28 days were each transferred to a separate well within flat-bottom, 96-well microplates with culture medium supplemented with 25 mM HEPES (Gibco, 15630080), to enable pH control outside of the incubator environment. Videotaping of CiliaBot motility was performed at 5 frames per second (FPS) using a Nikon TS2 microscope equipped with a Moment CMOS camera.

CiliaBot videos were analyzed using custom MATLAB programs that automatically extracted the 2D projections of the CiliaBots across the entire runtime via edge detection to identify areas in the video frames of high pixel contrast as a result of CiliaBots being noticeably darker than background, before filling out the space between detected edges to generate a binary mask representing the CiliaBot. From these projections, the XY-position of the centroid and major axis orientation were determined through blob analysis of the binary mask, and these metrics were further used to calculate the average translational velocity, average rotational velocity, and average path curvature (Figure 2A). A Savitzky-Golay smoothing filter was applied to the outputs of major axis orientation to reduce the influence of noise on rotational velocity calculations.

To further reveal the dynamic relationship between CiliaBot rotation and position, the “path-and-extent” visualization method was developed to transduce quantitative motility data into a single concise image. To enable this visualization, the XY-positions of the CiliaBot vertices were further obtained through blob analysis of the CiliaBots’ binary mask: for UniBots and rod-shaped AggreBots, the vertices of the major axis; for triangle-shaped AggreBots, the three centroid-vertex line segments; and for diamond-shaped AggreBots, the vertices of both the major and minor axes. These additional obtained values were used to depict the “path-and-extent” of a given CiliaBot’s motility pattern, superimposed on a freeze-frame of the video’s end. The “path” of the visualization stems from plotting the XY-position of the centroid over time. The “extent” stems from depicting the major axis, minor axis, and/or centroid-vertex line segments every 30 seconds of the video’s runtime. When used for hybrid AggreBots, regions of immotile cilia were visually identified during videotaping, with the identified vertex being highlighted with red “extent” to contrast the green “extent” denoting regions of motile cilia.

### Immunofluorescence staining and imaging

CiliaBots were fixed with 4% paraformaldehyde (PFA) for 1 hour at 4°C and washed with phosphate-buffered saline (PBS) with 0.1% Tween-20 (PBST). For immunofluorescence staining, CiliaBots were permeabilized with 1% Triton-X in PBS for 45 minutes, blocked with 1% bovine serum albumin (BSA) in PBS, incubated with the mouse anti-Acetylated-α-Tubulin (Ac-α-Tub) (Sigma-Aldrich, T6793) overnight at 4°C. Following a PBST wash, they were then incubated with donkey anti-mouse IgG (H+L) secondary antibody, Alexa Fluor 647 (ThermoFisher Scientific, A-31571), for 45 min at room temperature, and nuclear counterstained with DAPI (4′,6-diamidino-2-phenylindole) (ThermoFisher Scientific, D1306). Z-stack images of stained CiliaBots were captured on a Nikon A1 confocal microscope (*27*).

### Morphological cilia coverage analysis

From Z-stack images of CiliaBots stained with Acetylated-α-Tubulin and DAPI, 3 z-slices in or near the CiliaBot’s midsection were selected. Pixels of DAPI expression were used to generate a convex hull representing the tissue area, from which the centroid was calculated. From the calculated centroid, each z-slice was divided into 1-degree angular segments with an axis of rotation perpendicular to the image slice (Figure 1D). Finally, using a custom MATLAB program, the presence of Ac-α-Tub signal outside of the DAPI convex hull, limiting search only to signal found on the CiliaBot’s exterior surface, was evaluated in each angular segment, with a running tally being kept of which segments of Ac-α-Tub signal were above a threshold set to 10% of the maximum Ac-α-Tub signal intensity in the image. The count of angular segments containing Ac-α-Tub signal was then divided by 360 to calculate the morphological cilia coverage (*27, 33*).

### Functional cilia coverage analysis

AggreBots were transferred to a 1.5-mL Eppendorf tube and kept on ice. Ice-cold collagen type 1 (Advanced BioMatrix, 5225) was neutralized and added the Eppendorf tube at a concentration of 2 mg/mL, with 200 μL of this mixture of AggreBot and pre-gel collagen being transferred to a glass-bottom dish (Mattek, P35G-1.5-14-C). This dish was kept on ice for 10 minutes to allow the AggreBots to settle at the bottom. The Mattek dish was then incubated at 37°C for 10 minutes to allow collagen gelation before adding 1 mL of differentiation medium. Video recording of cilia beating was captured using an EVOS M7000 microscope with a 40X objective camera at 30 FPS.

Videos of collagen-embedded AggreBots were first processed with a video stabilization tool ((https://www.onlineconverter.com/stabilize-video), to eliminate motion within the video stemming from jitter. A Gaussian filter of 0.5 deviations was further applied to smooth any remaining artifacts. Following this pre-processing, the range of pixel intensities over the course of the captured video were calculated from each pixel and converted into a heatmap to identify regions of high motion activity, from which a custom MATLAB program identified the presence and size of the active cilia domain through the identification of high-intensity pixels in 360 angular segments along the surface of the AggreBots.

### AggreBot boundary integrity analysis

To assess boundary integrity, Aggrebots were formed from two CBBs stained with DiD (ThermoFisher Scientific, V22887) and DiO (Biotium, 60015) fluorescent dyes at 1 μM, respectively. These stained CBBs were aggregated on Day 1 and treated with either YAP activator XMU-MP-1 (Tocris Bioscience, 6482) at 10 μM, Rac1 inhibitor NSC23766 (Millipore Sigma, SML0952) at 50 μM, or both. Untreated CBBs were aggregated as a control group. 13 days following aggregation, Z-stack images of the resulting immature AggreBots were captured on a Zeiss LSM 700 confocal microscope, from which the middle image slice was selected. An image profile running the AggreBot’s major axis length was then extracted using a custom MATLAB program, and the pixel intensity relative to the maximum of both green and red channels was charted. The correlation coefficient between the green and pixel intensity pairs across the major axis length was then calculated.

## Supplementary Materials

**Supplementary Fig. S1**. Inability of mature CiliaBots to aggregate.

**Supplementary Fig. S2**. Aggregation success rate as function of CBB age.

**Supplementary Fig. S3**. Schematic depiction of process behind characterization of CiliaBot motility.

**Supplementary Movie S1**. Exterior cilia agitate CiliaBots, preventing stable contact and aggregation.

**Supplementary Movie S2**. Locomotion of a CBB_2_ AggreBot, depicting characteristic loop-de-loop behavior.

**Supplementary Movie S3**. Locomotion of a CBB_3_ AggreBot.

**Supplementary Movie S4**. Locomotion of a CBB_4_ AggreBot.

**Supplementary Movie S5**. Locomotion of a ^ND^CBB UniBot.

**Supplementary Movie S6**. Lack of motility from a ^PCD^CBB UniBot.

**Supplementary Movie S7**. Cilia activity of a collagen-embedded ^ND^CBB_1_^PCD^CBB_1_ AggreBot.

**Supplementary Movie S8**. Cilia activity of a collagen-embedded ^ND^CBB_2_^PCD^CBB_1_ AggreBot.

**Supplementary Movie S9**. Cilia activity of a collagen-embedded ^ND^CBB_1_ ^PCD^CBB_2_ AggreBot.

**Supplementary Movie S10**. Locomotion of a ^ND^CBB_2_ AggreBot.

**Supplementary Movie S11**. Locomotion of a ^ND^CBB ^PCD^CBB_1_ AggreBot, showcasing the decreased translational speed and increased path curvature brought about by the incorporation of ^PCD^CBBs.

**Supplementary Movie S12**. Locomotion of a ^ND^CBB_3_ AggreBot.

**Supplementary Movie S13**. Locomotion of a ^ND^CBB_1_ ^PCD^CBB_2_ AggreBot.

**Supplementary Movie S14**. Locomotion of a ^ND^CBB_4_ AggreBot.

**Supplementary Movie S15**. Locomotion of a chiral ^ND^CBB_2_ ^PCD^CBB_2_ AggreBot.

## Acknowledgments

We are grateful to Misti West and Garrett Struble for laboratory management, the Center for Biologic Imaging at the University of Pittsburgh and Dr. Adam Feinberg (Carnegie Mellon University) for access to imaging facilities and equipment, Dr. Patjanaporn Chalacheva (Carnegie Mellon University) for consulting on matters of data and statistical analysis, and Dr. Hongmei Mou (Massachusetts General Hospital and Harvard Medical School) for providing the 804G cell line. All figures were created in https://BioRender.com.

## Author contributions

Conceptualization: DB, XR

Methodology: DB, XZ, KG, SY, XR

Materials: SLB, AH

Investigation: DB, XZ, KG, SY, XR

Visualization: DB, ZG

Funding acquisition: DB, VW, ABF, XR

Project administration: XR

Supervision: XR

Writing – original draft: DB

Writing – review & editing: DB, SLB, AH, VW, ABF, XR

## Data and materials availability

All data, code, materials, and protocols used in the completion of this research will be made available upon request.

