## Supplementary Figures S1-3 for "AggreBots: configuring CiliaBots through guided, modular tissue aggregation"

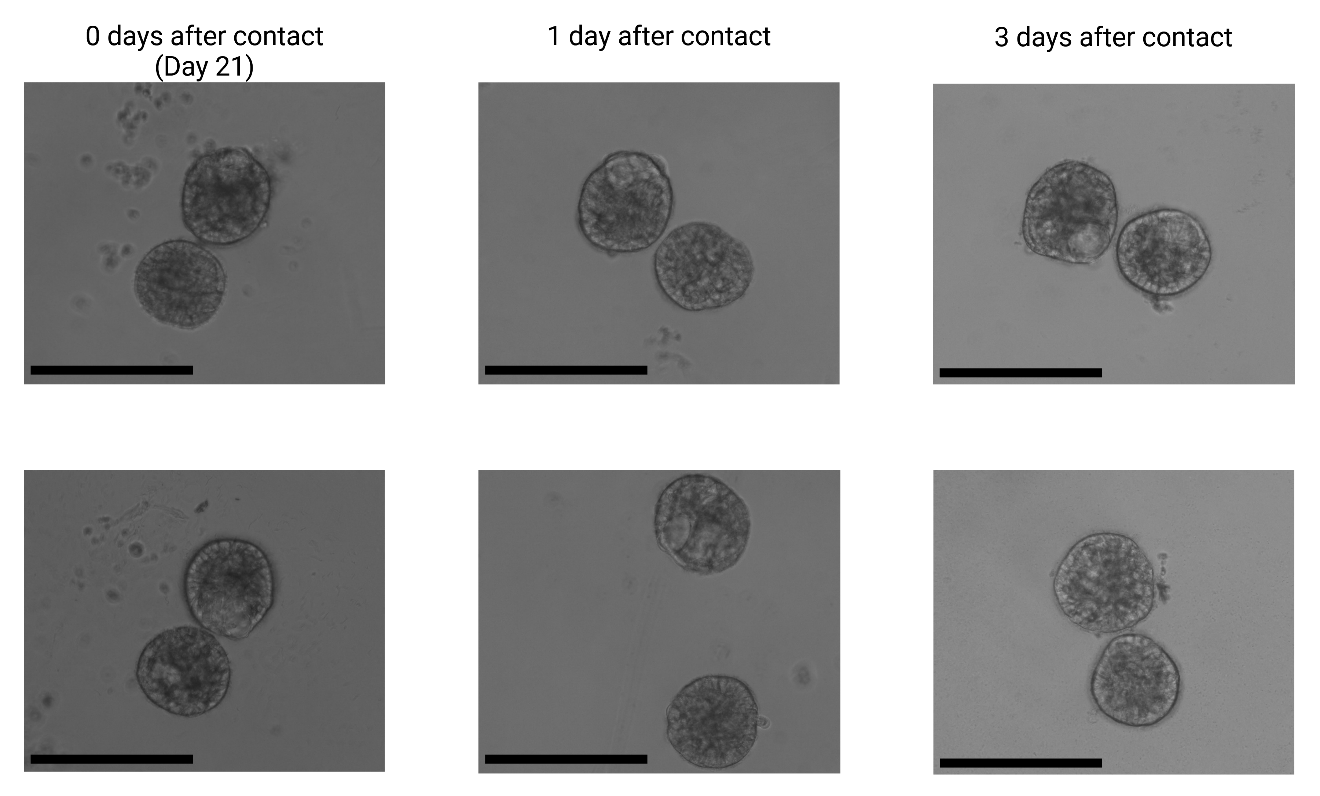

**Supplementary Fig. S1.** **Inability of mature CiliaBots to aggregate.** CiliaBots were brought into contact with one another on Day 21. Images were subsequently taken 1 and 3 days following initial contact, showcasing that mature CiliaBots appear to have no tendency to aggregate.

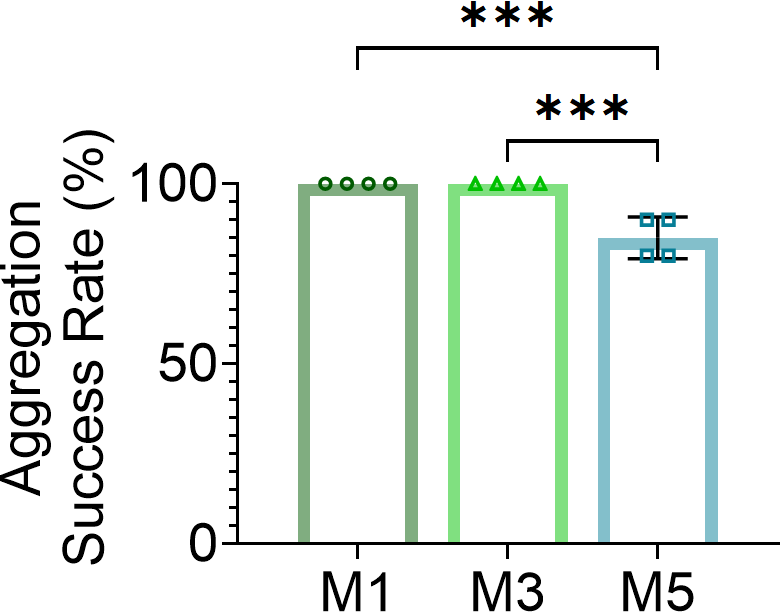

**Supplementary Fig. S2.** **Aggregation success rate as function of CBB age.** Aggregation success rates of each CBB age pre-contact group (M1, M3, M5). Each data point represents a separate experimental batch of CBBs.

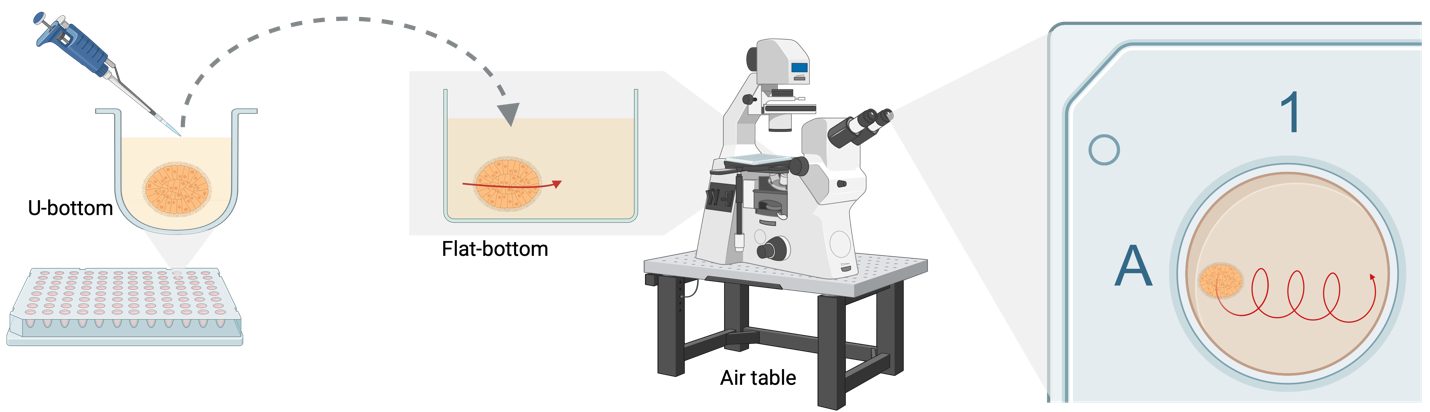

**Supplementary Fig. S3.** Schematic depiction of process behind characterization of CiliaBot motility.

**Supplementary Movie S1.** Exterior cilia agitate CiliaBots, preventing stable contact and aggregation.

**Supplementary Movie S2.** 30X Speed video depicting locomotion of a CBB_2_ AggreBot, alongside visualization of the traced loop-de-loop via the “spine-and-vertebrae” method.

**Supplementary Movie S3.** Locomotion of a CBB_3_ AggreBot.

**Supplementary Movie S4.** Locomotion of a CBB_4_ AggreBot.

**Supplementary Movie S5.** Locomotion of a ^H^CBB UniBot.

**Supplementary Movie S6.** Lack of motility from a ^PCD^CBB UniBot.

**Supplementary Movie S7.** Cilia activity of a collagen-embedded ^H^CBB_1_^PCD^CBB_1_ AggreBot.

**Supplementary Movie S8.** Cilia activity of a collagen-embedded ^H^CBB_2_^PCD^CBB_1_ AggreBot.

**Supplementary Movie S9.** Cilia activity of a collagen-embedded ^H^CBB_1_^PCD^CBB_2_ AggreBot.

**Supplementary Movie S10.** Locomotion of a ^H^CBB_2_ AggreBot.

**Supplementary Movie S11.** Locomotion of a ^H^CBB_1_^PCD^CBB_1_ AggreBot, showcasing the decreased translational speed and increased path curvature brought about by the incorporation of ^PCD^CBBs, with ^PCD^CBB marked with red “vertebrae”.

**Supplementary Movie S12.** Locomotion of a ^H^CBB_3_ AggreBot.

**Supplementary Movie S13.** Locomotion of a ^H^CBB_1_^PCD^CBB_2_ AggreBot.

**Supplementary Movie S14.** Locomotion of a ^H^CBB_4_ AggreBot.

**Supplementary Movie S15.** Locomotion of a chiral ^H^CBB_2_^PCD^CBB_2_ AggreBot.
